# Single-cell proteomics of Arabidopsis leaf mesophyll identifies drought stress-related proteins

**DOI:** 10.1101/2024.07.01.601628

**Authors:** James M. Fulcher, Pranav Dawar, Vimal Kumar Balasubramanian, Sarah M. Williams, Kyle D. Nadeau, Tanya E. Winkler, Lye Meng Markillie, Hugh D. Mitchell, Amir H. Ahkami, Ljiljana Paša-Tolić, Ying Zhu

## Abstract

The application of single-cell omics tools to biological systems can provide unique insights into diverse cellular populations and their heterogeneous responses to internal and external perturbations. Thus far, most single-cell studies in plant systems have been limited to RNA-sequencing approaches, which only provide indirect readouts of cellular functions. Here, we present a single-cell proteomics workflow for plant cells that integrates tape-sandwich protoplasting, piezoelectric cell sorting, nanoPOTS sample preparation, and FAIMS-based MS data acquisition method for label-free single-cell proteomics analysis of Arabidopsis leaf mesophyll cells. From a single leaf protoplast, over 3,000 proteins were quantified with high precision. The workflow is demonstrated to identify stress associated changes in protein abundance by analyzing >80 protoplasts from well-watered and water-deficit stressed plants. Additionally, we describe a new approach for constructing covarying protein networks at the single-cell level and demonstrate how single-cell protein covariation analysis can reveal previously unrecognized protein functions while also capturing stress-induced changes in protein-protein dynamics.

**Highlights:**

- This study describes the first application of scProteomics to study abiotic stress-induced proteome regulation in leaf mesophyll cells
- From a single Arabidopsis mesophyll protoplast, ∼2800 proteins on average were quantified with high precision using label free scProteomics
- scProteomics of water-deficit Arabidopsis leaves revealed known and novel proteins involved in drought response
- Single-protoplast proteomics revealed water deficit-induced changes in protein covariation networks independent of changes in abundance

## Introduction

Abiotic stresses have led to decreases in agricultural productivity, and these issues are anticipated to increase in the future^1^. Whole tissue (“bulk”) -omic techniques have frequently been used to identify abiotic stress responsive genes that are amenable to genome editing towards producing more resilient and productive crops ^2–4^. Recently, the introduction of single-cell omics technologies in plant systems has revealed the heterogeneity and spatiotemporal regulation of plant cells in various contexts, including abiotic stress and early stages of development ^5–9^. The majority of these approaches have relied on next-generation sequencing, providing genomic, epigenomic, and transcriptomic measurements^6^. Of these, single-cell RNA sequencing (scRNAseq) of mRNA has been more often utilized due to its high-throughput capabilities and accessibility. Unfortunately, the dynamic (and sometimes stochastic) nature of mRNA expression means it is an imprecise predictor of protein expression levels^10,11^. This is further exacerbated at the single-cell level compared to bulk tissue level^10^. To this end, recent emerging single-cell omics platforms have focused on measuring protein abundance at a global level through mass spectrometry^12^. Although many single-cell proteomics (scProteomics) studies have been performed in mammalian systems, cell-type specific proteomics and scProteomics in plants has been rarely reported^13–16^. This is primarily because plants do not produce free-moving single cells during their life cycle, apart from pollen grains and spores. Hence, one bottleneck in plant single-cell proteomics has been obtaining cells in sufficient quantities and quality. Encouragingly, a recent study demonstrated the feasibility of tandem-mass tag (TMT)-based scProteomics on plant samples following the SCoPE protocol^17,18^. With a focus on distinguishing two root cell types (cortex and endodermis), the authors indicated cell-type specific protein markers could be obtained ^16^. However, one limitation of this approach is the reliance on a large isobaric carrier channel (∼140x) which leads to fewer ions for quantification from single-cells and can therefore have negative impacts on protein measurement dynamic range and quantitative accuracy^19,20^.

Here, we developed and evaluated a label-free scProteomics workflow to study drought-induced cell response using *Arabidopsis thaliana* leaf mesophyll cells.

Mesophyll cells were considered as an ideal model due to their large size, high abundance in leaf tissue, and their unique protein responses under water stress^21^. The workflow for analyzing these cells integrates tape-sandwich protoplasting^22^, piezoelectric cell sorting (cellenONE), nanodroplet Processing in One Pot for Trace Samples (nanoPOTS)^23^, and high field asymmetric waveform ion mobility spectrometry (FAIMS)-enhanced data acquisition^24^. Because cell-wall lacking protoplasts are very fragile, it is challenging to survive in high-pressure sheath-flow-based sorting system (e.g., flow cytometry)^25^. We show the cellenONE low-pressure cell sorting system can greatly reduce protoplast membrane breakage. The cellenONE imaging capabilities could also be used to select intact protoplasts and avoid poor-quality fragmented protoplasts that may confound downstream analysis. With the FAIMS-enhanced scProteomics acquisition method, 2,873 proteins were identified on average from well-watered and water-deficit (WD) stressed single Arabidopsis leaf mesophyll cells. Lastly, these scProteomics datasets allowed us to construct covarying protein networks to identify novel protein functions and protein-protein interactions under WD stress.

## Experimental Procedures

### Materials

Deionized water (18.2 MΩ) was purified using a Barnstead Nanopure Infinity system (Los Angeles, CA, USA). N-dodecyl-β-D-maltoside (DDM), iodoacetamide (IAA), ammonium bicarbonate (ABC), and formic acid (FA) were obtained from Sigma (St. Louis, MO, USA). Trypsin (Promega, Madison, WI, USA) and Lys-C (Wako, Japan) were dissolved in 50 mM ABC before usage. Dithiothreitol (DTT, No-Weigh format), acetonitrile (ACN) with 0.1% FA, and water with 0.1% FA (MS grade) were purchased from Thermo Fisher Scientific (Waltham, MA, USA). Jack’s professional fertilizer was acquired from Jr Peters Inc., PA, USA and SunGro soil was acquired from SunGro, OR, USA. CELLULYSIN Cellulase was acquired from Millipore Sigma and macerozyme R10 was acquired from Fischer Scientific, USA. Polyimide Kapton tape was acquired from 3M (Maplewood, MN, USA).

### Fabrication of nanoPOTS chip

The nanoPOTS chips were fabricated using standard photolithography, wet etching, and silanization as described previously^23^. Each chip contained 48 (4 x12) nanowells with a well diameter of 1.2 mm and an inter-well distance of 4.5 mm. Chip fabrication utilized a 25 mm x 75 mm glass slide pre-coated with chromium and photoresist (Telic Company, Valencia, USA). After photoresist exposure, development, and chromium etching (Transene Company, Inc.), the exposed region was etched to a depth of ∼5 μm with buffered hydrofluoric acid. The freshly etched slide was dried by heating it at 120 °C for 1 h and then treated with oxygen plasma for 3 min (AP-300, Nordson March, Concord, USA). 2% (v/v) heptadecafluoro-1,1,2,2-tetrahydrodecyl-dimethylchlorosilane (PFDS, Gelest, Germany) in 2,2,4-trimethylpentane was applied onto the chip surface and incubated for 30 min to allow for silanization. The remaining chromium covering the wells was removed with chromium etchant, leaving elevated hydrophilic nanowells surrounded by a hydrophobic surface. Polyetheretherketone (PEEK) chip covers were machined as chip covers to provide sealing and protection of the droplets from evaporation during sample processing. Chips were wrapped in aluminum foil for long-term storage and intermediate steps during sample preparation.

### Plant growth conditions and water-deficit treatment

The *A. thaliana* (Columbia-0 ecotype) seeds were germinated on SunGro soil and kept in a reach-in growth chamber maintained at a temperature of 22 °C /18 °C (day/night), a light intensity of 150 μmol/m^2^/s and a photoperiod of 16 h/18 h (day/night). After four days, the germinated seedlings were transplanted (two seedlings per individual pot (4” × 4” × 9.5”). The Arabidopsis plants were fertilized once weekly using 100 ppm of Jack’s professional fertilizer. After two weeks of seedling growth, half of the plants were exposed to water deficit (WD) stress by reducing the watering levels to 50% of initial levels for the next two days, followed by stop-watering until the soil water content (SWC) arrived to 50% SWC (moderate WD, referred to as WD-50%) and 30% SWC (severe WD, referred to as WD-30%)^26^. SWC was calculated by measuring the dry soil weight and water weight needed for full saturation of soil at the start of the experiment. Control and WD stress-induced pot weights were measured daily to calculate soil weights required to achieve 50% and 30% SWC (supplemental Fig. S1A). It took approximately seven days of drying to achieve 50% SWC and another week to reach 30% SWC. While the moderate WD plants were harvested the next day after reaching 50% SWC, the plants at 30% SWC were maintained at that state for an additional three days to induce severe WD stress symptoms before phenotyping and harvesting were performed (supplemental Fig. S1B). The rosette leaf samples were collected between 9-10 AM from control and WD stressed plants and immediately used for protoplast isolation. For plant phenotyping, a separate set of plants was harvested to record leaf and root fresh weights and dry weights (after drying biomass at 60°C for five days). WD-50% plants showed no significant changes in leaf fresh weights and only a moderate reduction in leaf dry weight (supplemental Fig. S1C); however, the WD-30% plants had a significant reduction in leaf fresh and dry weights (supplemental Fig. S1D).

### Protoplast isolation

We followed the *Arabidopsis* tape-sandwich protocol to isolate the leaf protoplast enriched with palisade mesophyll cells^22^. Briefly, control and WD-stressed *Arabidopsis* rosette leaves were placed on 3M tape, and abaxial cell layers from leaf were peeled off by gently removing the tape, thereby exposing the adaxial side with enriched palisade mesophyll cells. The exposed leaf (n=6) from control and WD stress conditions was placed in a petri plate, and 20 ml of protoplasting enzyme solution was added. The enzyme solution contained 1% cellulase (CELLULYSIN^®^) and 0.25% macerozyme R10 dissolved in a buffer made of 0.4M mannitol, 20 mM potassium chloride, 20 mM 2-(N-morpholino)ethanesulfonic acid (MES) buffer and adjusted to pH 5.7. The enzyme solution was heated at 55°C for 10 mins to dissolve the enzymes into the buffer followed by cooling the solution on ice for 5 minutes. Following that, 10 mM calcium chloride was added to the enzyme solution, and it was used for protoplast isolation. The leaf tissue was incubated with the enzyme solution for 45 minutes and kept on a rotatory shaker maintained at 150rpm at 25°C in the dark to allow the enzymes to digest the cell walls to liberate the protoplasts. Three batches of protoplast isolation were performed from control and WD-stressed leaf tissue to yield sufficient protoplasts for single-cell proteomics workflow. After enzymatic digestion, the protoplast solution was carefully transferred to 50ml falcon tubes and centrifuged at 13 G at room temperature using an IEC Centra MP4R refrigerated centrifuge. The protoplast pellet was gently washed two times using the protoplast isolation buffer without mannitol and filtered using a 0.45 μm filter to remove the cell debris. The filtered leaf protoplasts were counted using hemocytometer and immediately sorted using the cellenONE.

### Sorting of protoplasts with cellenONE

A cellenONE instrument equipped with a glass piezo capillary (P-20-CL) for aspiration and dispensing was utilized for single-cell isolation. Sorting parameters included a pulse length of 51 μs, a nozzle voltage of 87 V, a frequency of 500 Hz, an LED delay of 200 μs, and an LED pulse of 3 μs. The slide stage was operated at dew-point control mode to reduce droplet evaporation and keep protoplasts at low temperature after sorting (∼2-4 °C). Protoplasts were isolated based on their size, circularity, and elongation to exclude fragmented protoplasts, doublets, or other cell debris. Autofluorescence (excitation of 625nm and emission of 670nm) images were also collected during sorting. Specific criteria for selection included cell diameters of 40 to 70 μm, maximum circularity of 2, and maximum elongation of 2.7. All protoplasts were sorted based on bright field images in real time. To perform the transferring identifications based on FAIMS filtering (TIFF) methodology for scProteomics^24^, a library chip was also prepared containing 20 cells per nanowell. After sorting, all chips were wrapped in aluminum foil before being snap-frozen and stored at -80°C.

### nanoPOTS-based proteomic sample preparation

For scProteomics with nanoPOTS, extraction was accomplished by dispensing 150 nL of extraction buffer containing 50 mM ABC, 0.1% DDM, 0.5 x diluted PBS, and 2 mM DTT and incubating the chip at 60°C for 60 min. Denatured and reduced proteins were alkylated through the addition of 50 nL 15 mM IAA before incubation for 30 min in darkness at room temperature. Alkylated proteins were then digested by adding 50 nL 50 mM ABC with 0.01 ng/nL of Lys-C and 0.04 ng/nL of trypsin and incubating at 37°C overnight. The digestion reaction was then quenched by adding 50 nL of 5% formic acid before drying the chip under vacuum at room temperature. All chips were stored in at -20°C until LC-MS analysis.

### LC-MS/MS

We employed an in-house assembled nanoPOTS autosampler for LC-MS/MS analysis. The autosampler contains a custom packed SPE column (100 μm i.d., 4 cm, 5 μm particle size, 300 Å pore size C18 material, Phenomenex) and an analytical LC column (50 μm i.d., 25 cm long, 1.7 μm particle size, 190 Å pore size, C18 BEH material, Waters) with a self-pack picofrit (cat. no. PF360-50-10-N-5, New Objective, Littleton, MA). The analytical column was heated to 50 °C using AgileSleeve column heater (Analytical Sales and services, Inc., Flanders, NJ). Briefly, samples were dissolved with Buffer A (0.1% formic acid in water) on the chip, then trapped on the SPE column for 5 min. After washing the peptides, samples were eluted at 100 nL/min and separated using a 30 min gradient from 8% to 35% Buffer B (0.1% formic acid in acetonitrile).

An Orbitrap Lumos or Eclipse Tribrid MS (ThermoFisher Scientific) with FAIMSpro, operated in data-dependent acquisition mode, was used for all analyses. Source settings included a spray voltage of 2,400 V, ion transfer tube temperature of 250°C, and carrier gas flow of 4.6 L/min. For individual sampes^24^, ionized peptides were fractionated by the FAIMS interface using internal compensation voltage stepping (-45, -60, and -75 V) with a total cycle time of 0.8 s per compensation voltage. Fractionated ions within a mass range 350-1600 m/z were acquired at 120,000 resolution with a max injection time of 500 ms, AGC target of 1E6, RF lens of 30%. Tandem mass spectra were collected in the ion trap with an AGC target of 20,000, a “rapid” ion trap scan rate, an isolation window of 1.4 *m/z*, a maximum injection time of 120 ms, and a HCD collision energy of 30%. For the TIFF library samples, a single compensation voltage was used for each LC-MS run with slight modifications to the above method where cycle time was increased to 2 s and maximum injection time was set to 118 ms. Precursor ions with a minimum intensity of 1E4 were selected for fragmentation by 30% HCD and scanned in an ion trap with an AGC of 2E4 and an IT of 150 ms.

### Database searching and data processing

All proteomic data raw files were processed by FragPipe^27^ (version 21.0) and searched against the *Arabidopsis thaliana* UniProt protein sequence database (UP000006548 containing 41,594 forward and reversed entries accessed 03/24) including decoy sequences and common protein contaminants. Search settings included a precursor mass tolerance of +/- 20 ppm, fragment mass tolerance of +/- 0.5 Da, deisotoping, trypsin as the enzyme with N-terminal semi-tryptic specificity, carbamidomethylation as a fixed modification, and several variable modifications (oxidation of methionine and N-terminal acetylation). Protein and peptide identifications were filtered to a false discovery rate of less than 0.01 within FragPipe. For the TIFF methodology, IonQuant match-between-runs (MBR) and MaxLFQ were set to “TRUE” and library MS datasets were assigned as such during the data import step. An MBR FDR of 0.05 at ion level was used to reduce false matching.

FragPipe result files were then imported into RStudio (2024.04.1 Build 748) for downstream analysis in the R environment (version 4.4.0). All datasets were median normalized at the peptide level using the “median_normalization” function from the “proDA” R package before rolling up to MaxLFQ protein abundances using the “fast_MaxLFQ” function from the “iq” R package^28–30^. For cases where complete data was needed, imputation was performed with k-nearest neighbors imputation (n = 10 neighbors) on features with < 10% missing values. Gene ontology analysis was performed with the gprofiler2 R package^31^, while gaussian mixture modeling (GMM) clustering was accomplished with the Mclust R package^32^. Note for GO enrichment analysis we used a more stringent domain scope instead of the full *Arabidopsis* proteome. This domain scope consisted of the 3,587 proteins identified within our study. The intensity Based Absolute Quantification (iBAQ) values were calculated by summing all peptide intensities for a given protein and dividing it by the number of theoretically observable tryptic peptides between 6 and 50 amino acids in length^33^. These values were then converted to mol fraction (also referred to as relative iBAQ or riBAQ) by summing the total iBAQ signal for each sample and dividing each protein by this total, excluding contaminant proteins^34^. Finally, mol fractions were converted into copy number using previously calculated cellular protein copy number (25.8 billion) for *Arabidopsis* leaf mesophyll cells^21^.

### Construction of protein covariation networks

For the generation of the consensus protein covariation network, we first identified the set of clusters (k) from GMM clustering that provided the most accurate representation of the network relative to known gene ontologies without significant cluster splitting or other artifacts. Briefly, this was accomplished by performing GO enrichment analysis on each cluster from a given GMM model with k number of clusters. For each statistically significant term (FDR corrected hypergeometric test p-values of < 0.05) associated with a given cluster, we calculated a weighted (*β*) F-score (F) defined as the harmonic mean of the precision and recall (Equation 1). Note that precision is defined as the proportion of proteins in a cluster overlapping with the set of proteins from a GO term relative to the full set of proteins within the cluster. On the other hand, recall represents the proportion of functionally annotated proteins that intersect with a cluster relative to the entire set of proteins within that GO term. Because scProteomics is typically more limited in coverage of the proteome, we rationalized that recall is going to be less predictive of successful clustering since the absence of proteins is more likely to be attributable to low abundance and therefore not measurable during LC-MS analysis. For this reason, we selected a *β* of 0.5 to weigh precision at twice the weight of recall.

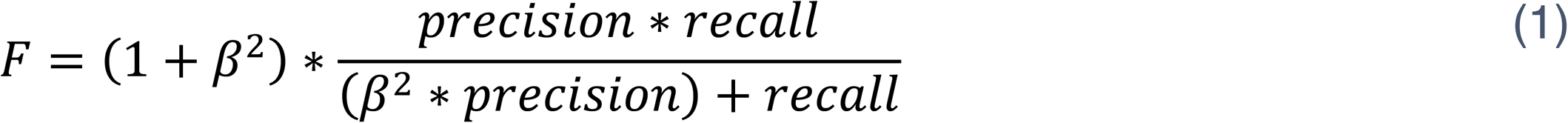

Clusters that are found to have an enriched GO term are referred to as kx below. As each cluster may contain multiple GO terms associated with it, only the term with the highest F-score was used in the calculations below. Each cluster’s maximum F-score was summed together provided they belong to unique, non-redundant statistically significant GO terms. Two correction factors are then used to provide the final “GO-Score” associated with each k tested (Equation 2).

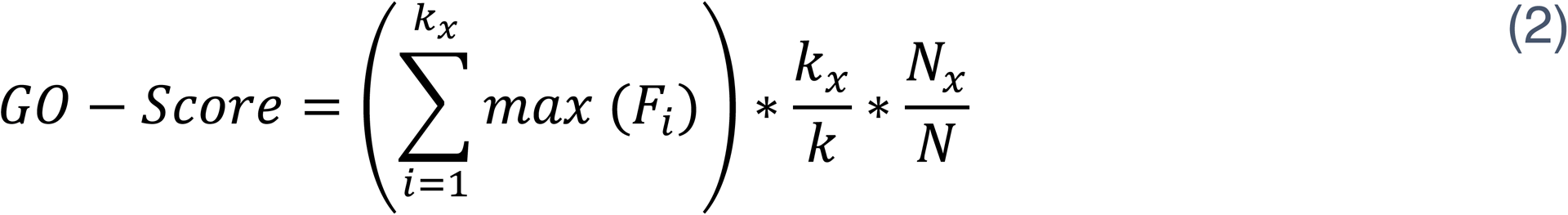

The first correction factor scales as k clusters have proportional increases in kx, and therefore penalizes increasing numbers of k when kx does not increase. The second correction factor provides greater weight to k clusters that contain more GO annotated proteins (Nx) relative to the total proteins input into the GMM (N). The highest GO-Score, therefore, was used to identify the best k clusters after GMM to be used in downstream analysis. To visualize the network, pairwise protein-protein correlations within or across each kx were averaged. Euclidean distance and hierarchical clustering were then used to identify the order of clusters within the heatmap. Proteins within each cluster were further organized from lowest to highest by cluster uncertainty values output from the GMM. Note that uncertainty in this context refers to the statistical confidence of a protein’s assignment to a particular cluster for a given k. This final arrangement was then visualized by the ComplexHeatmap R package and a summary table was produced with the “gt” R package ^35,36^. Associated code and functions are provided at the Github repository (https://github.com/Cajun-data/nanoPOTS_Arabidopsis).

### Experimental Design and Statistical Rationale

For technology evaluation with different number of *Arabidopsis* protoplasts, experiments were performed with 20 library protoplasts (n = 6), 10 pooled protoplasts (n = 4), 3 pooled protoplasts (n = 4), and single protoplasts (n = 8). Note the library samples were necessary for IonQuant MBR using the TIFF methodology^24^. To sample across biological replicates, protoplasts were derived from six *Arabidopsis* plants (leaf specifically) grown with no water deficit stress pooled together during protoplasting. For the comparison of WD stress and control *Arabidopsis* protoplasts, the Ctrl-30%, Ctrl-50%, and WD-50% groups had n = 6 library samples consisting of 20 pooled protoplasts (n = 18 total). The total number of single protoplasts for each group were as follows: n = 28 Ctrl-50%, n = 30 for Ctrl-30%, n = 29 for WD-50%, and n = 30 for WD-30%. As before, in each condition, protoplasts were derived from six *Arabidopsis* plant leaf samples to sample across biological replicates.

A QC filtering step was undertaken to remove fragmented protoplasts which were found to add confounding effects to downstream analysis that could not be corrected by quantile normalization or batch effect correction. Specifically, fragmented protoplasts were identified using kmeans clustering on the first two principal components after principal component analysis (optimal k - which in this case was 2 - was chosen using within-cluster-sum of squared errors). Further confirmation of fragmented protoplasts was determined using images taken by the cellenONE instrument during sorting. In total, 30 fragmented protoplasts were clustered and removed, leaving 87 high-quality intact protoplasts across the four groups. This number of samples still provided sufficient statistical power to perform differential abundance and protein covariation analysis across protoplasts. Importantly, we assessed the quality, data completeness, and sensitivity of all single-protoplast samples at the protein level (3,587 unique identifications) and peptide level (30,018 unique identifications) as recommended by Vanderaa and Gatto (supplemental Fig. S2)^37^.

## Results

### nanoPOTS-based label free scProteomics workflow enables deep coverage in single mesophyll protoplasts

To enrich the Arabidopsis leaf mesophyll cell type, we adopted the tape-sandwich approach to remove abaxial and adaxial epidermal cell layers (Fig. 1A and supplemental Fig. S3) for downstream protoplasting^22^. Our initial attempt to use flow cytometry to sort protoplasts provided us negative results (0 cells observed from 25 total sorting events), when examining under the microscope. Presumably, these fragile protoplasts were lysed under the high-pressure sheath flow before reaching the sorting nozzle. This is not entirely unexpected, as the protoplasting process involves removing the cell wall which is known to make the cells fragile and prone to rupture during centrifugation steps or pipetting^25^. In contrast, the cellenONE sorts single cells without requiring high-pressure sheath flow. Instead, single cells are directly dispensed from the nozzle tip. Additionally, cells in the nozzle tip are imaged before dispensing, which can be used to validate cellular membrane integrity and avoid co-isolation of other cellular debris before dispensing of the protoplast. The cellenONE allowed us to achieve a much higher success rate for protoplast sorting, however ∼20% fragmented protoplasts were still observed during sorting, which can be attributed to the leaf state (control or stressed) and protoplasting procedures.

**Fig. 1.**
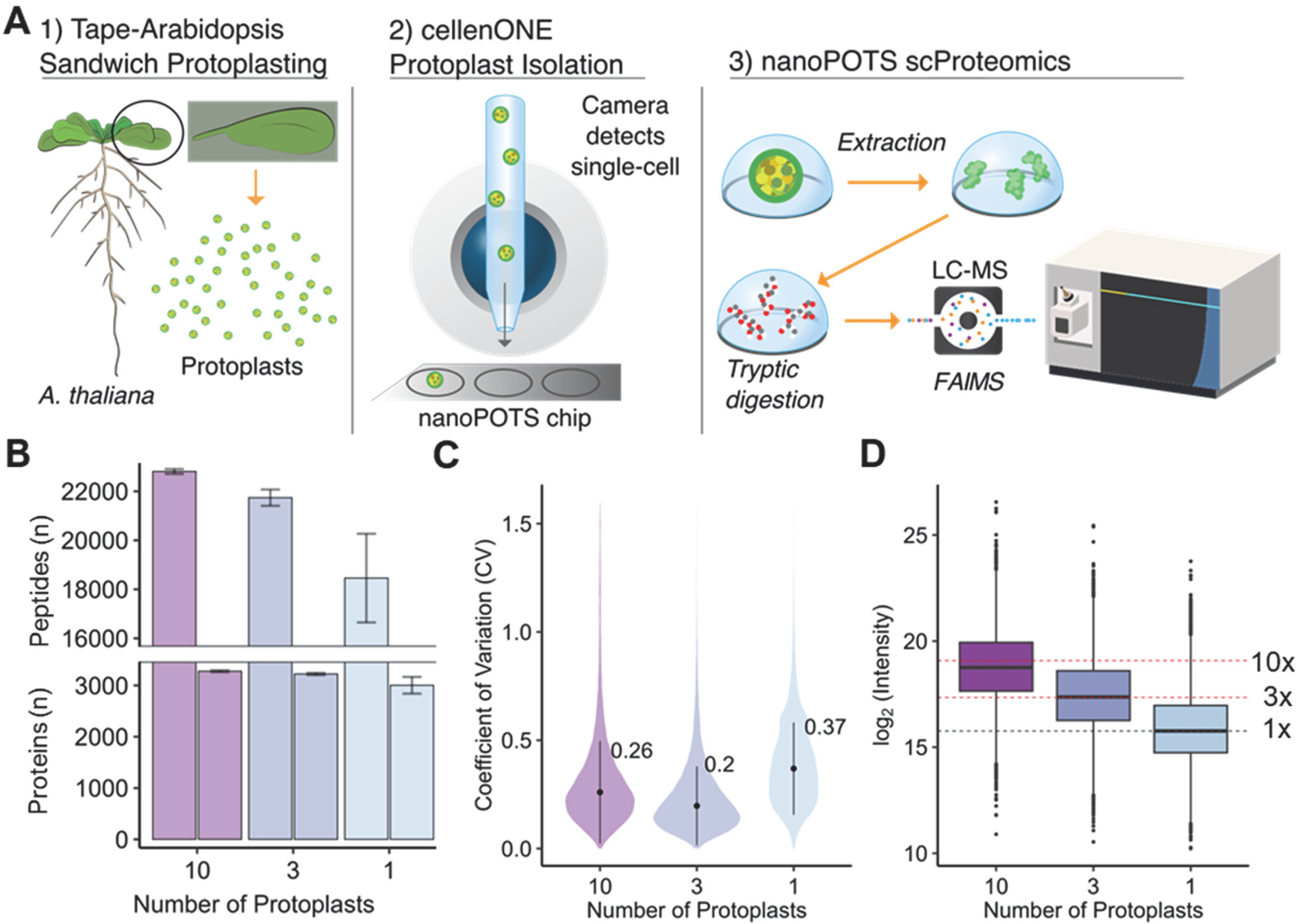
Proteome coverages and quantitation accuracy/precision based on nanoPOTS-based label-free scProteomics. **(A)** Overview of the nanoPOTS-based label-free scProteomic approach (**B**) Total peptide and protein identifications across 10 (n = 4), 3 (n = 4), and 1 (n = 8) protoplast. Error bars represent +/- SD. (**C**) Peptide level coefficients of variation for 10, 3, and 1 protoplast. (**D**) Log_2_ transformed protein intensity distributions for 10, 3, and 1 protoplast. Black dashed line indicates median of a single protoplast. Red dashed lines indicate theoretical 3-fold and 10-fold larger median abundances.

As a first step, we sought to determine the proteome depth and quantitation accuracy/precision of nanoPOTS-based scProteomics approach. We analyzed pools of 10, 3, and 1 protoplast with multiple replicates for each number (Fig.1B). As expected, a general trend of decreasing peptide and protein identifications was found from 10 mesophyll protoplasts to single-protoplasts. Notably, 18,458 peptides corresponding to 3,003 proteins on average were identified from a single protoplast and 22,812 peptides corresponding to 3,278 proteins were identified from 10 protoplasts. This great depth can be attributed to the large sizes of Arabidopsis mesophyll protoplasts with diameters typically ranging from 30 to 50 μm^38,39^. Indeed, protein content of Arabidopsis leaf cells has been estimated at ∼1.7 ng per cell, compared to the ∼200 pg in HeLa cells^21,40^.

Measurement precision (as determined by peptide-level coefficient of variation, CV) was also found to be within ranges expected for label free quantitative (LFQ) analysis (Fig. 1C). A trend of increasing CV was also noted from 10 to 1 protoplast, with 10 protoplasts having a median CV 0.26 and single protoplasts a median CV of 0.37, which indicates the increasing biological heterogeneity of the single-cell samples. Although the exact protein input is not known given the variance in protoplast size, comparison of the overall distributions of protein abundances across the three input levels (10, 3, and 1 protoplast) reveals the medians of these distributions approximates theoretical ratios (Fig. 1D).

We also determined the Pearson correlations across all inputs at the protein-level and found most Pearson correlations were > 0.9 with only two single-protoplast samples being outliers that correlated very strongly with each other (supplemental Fig. S4).

Based on cellenONE-generated images of each sorted protoplast, we noted a more irregular appearance of the two outlier protoplasts compared to all other single-protoplast samples, which indicates breakage of the plasma membrane and loss of cellular contents (supplemental Fig. S5). While most fragmented protoplasts were excluded during cellenONE sorting based on circularity and size, a small proportion retained features that mimic intact protoplasts and were selected for sorting despite having appropriate thresholds in place. In any case, these fragmented protoplasts can be gated out for proteomics analysis or identified based on their proteome abundance profiles, which facilitates their removal during downstream analysis (*vide infra*).

### Protein signatures from single Arabidopsis mesophyll protoplasts in the context of water-deficit stress

Having established the baseline characteristics of nanoPOTS scProteomics for plant cells, we next applied it to study how leaf mesophyll cells responds to WD stress at the single-cell level. Two levels of WD stress were tested, moderate (50% SWC, referred to as WD-50%) and severe (30% SWC, referred to as WD-30%) along with associated non-WD control plants (referred to as Ctrl-50% and Ctrl-30%). In total, 117 protoplasts (approximately 30 from each group) were analyzed with nanoPOTS scProteomics. Initial PCA analysis with kmeans clustering (k = 2) revealed confounding effects across the sample groups (supplemental Fig. S6 and supplemental Fig. S7A). Protoplast identification depth was markedly lower in one of the two clusters (supplemental Fig. S7B), suggestive of protoplast fragmentation which was confirmed visually with images from the cellenONE (supplemental Fig. S7C). We therefore implemented a quality control filtering step to remove these putative fragmented protoplasts, providing 87 high quality protoplasts (or 74% of the total protoplasts analyzed) for downstream analysis. In agreement with our initial scProteomic analysis of mesophyll protoplasts, an average of 2,873 proteins were identified across these 87 protoplasts (Fig. 2A).

**Fig. 2.**
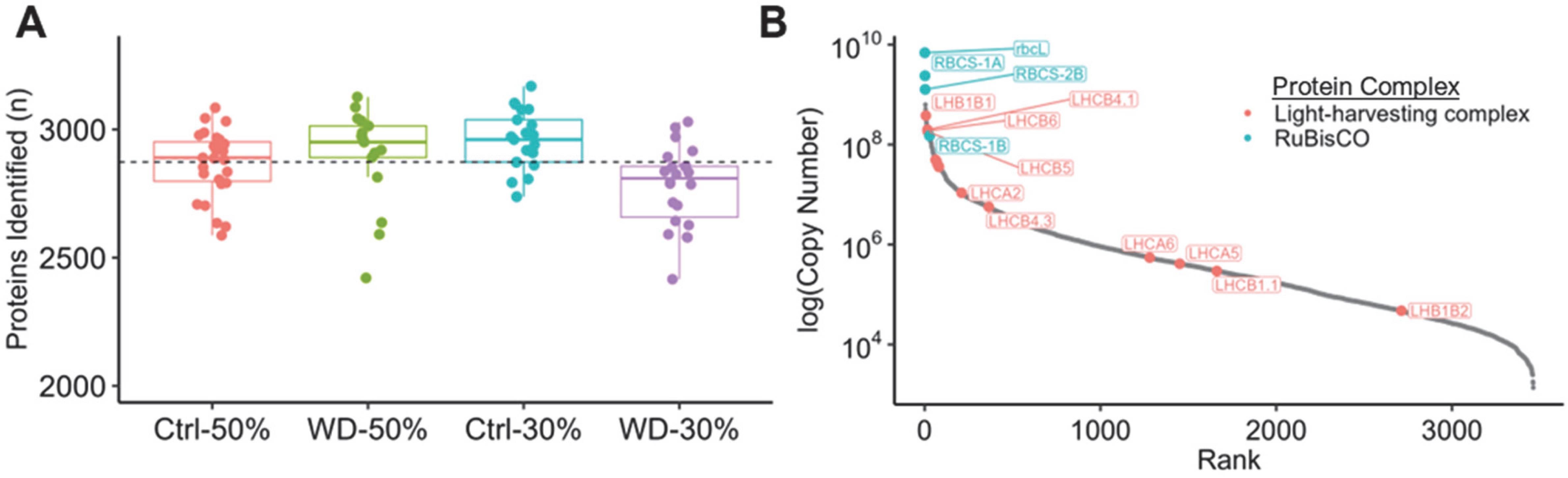
Initial characterization of water deficit stressed and well-watered mesophyll protoplasts. **(A)** The total number of proteins identified across control (Ctrl-50% and Ctrl-30%), moderate (WD-50%), and severe (WD-30%) water deficit protoplasts. Black dashed line indicates average number of protein identifications across all protoplasts (2,873). (**B**) Protein copy number abundances highlighting RuBisCo (blue color) and light-harvesting complex proteins (magenta color).

Conversion of protein LFQ intensities to copy numbers provided the absolute abundance of 3,462 proteins in our control sample protoplasts (n = 48, Fig. 2B and supplemental table S1). Two complexes were particularly overrepresented, Ribulose-1,5-Bisphosphate Carboxylase/oxygenase (RuBisCo) and photosystem I/II light-harvesting complexes (LHC), aligning with the expected function of mesophyll protoplasts being the major site of photosynthesis (Fig. 2B)^41^. Four out of five RuBisCo proteins and seventeen out of twenty LHC proteins were quantified, and the top three most abundant proteins were all RuBisCo subunits, which notably is also considered to be one of the most abundant proteins on Earth^41^. Indeed, our data suggests RuBisCo subunits alone account for ∼10 billion protein copies per single protoplast. The unidentified RuBisCo protein (RBCS-3B) was not attributed to low abundance or stochastic sampling during LC-MS analysis, but instead because it shares 97% sequence similarity RBCS-2B and hence could not be distinguished from RBCS-2B.

Interestingly, the LHC for photosystem I has four core proteins (LHCA1-4) conserved across higher plants as well as two additional LHC-type proteins (LHCA5 and LHCA6) that are thought to be differentially regulated and expressed at sub-stoichiometric amounts^42^. Our results support this observation and show that on average there are ∼78-fold fewer copies of LHCA5 and LHCA6 compared to the four core LHC photosystem I proteins (supplemental Fig. S8). As further validation of our mesophyll protoplast isolation process and quantitative accuracy, we compared the protein identification overlap and copy number correlation with a recently published shotgun-proteomic analysis of bulk Arabidopsis leaf tissue^21^. The protein identifications overlapped with 84% of the proteins identified by Heinemann et al. in bulk control leaf tissue and our study extended on their results by providing copy number estimates for an additional 2,270 proteins at single-protoplast resolution (supplemental Fig. S9A and supplemental table S1)^21^. Furthermore, among overlapping protein identifications we found our experimental copy number estimates to be strongly correlated (R = 0.79) with these prior results (supplemental Fig. S9B)^21^.

Principal component analysis (PCA) of all protoplasts suggests the effect of water deficiency is a major source of variance as the greatest separation between groups was found along the first principal component with 19.3% of the total variance (Fig. 3A).

**Fig. 3.**
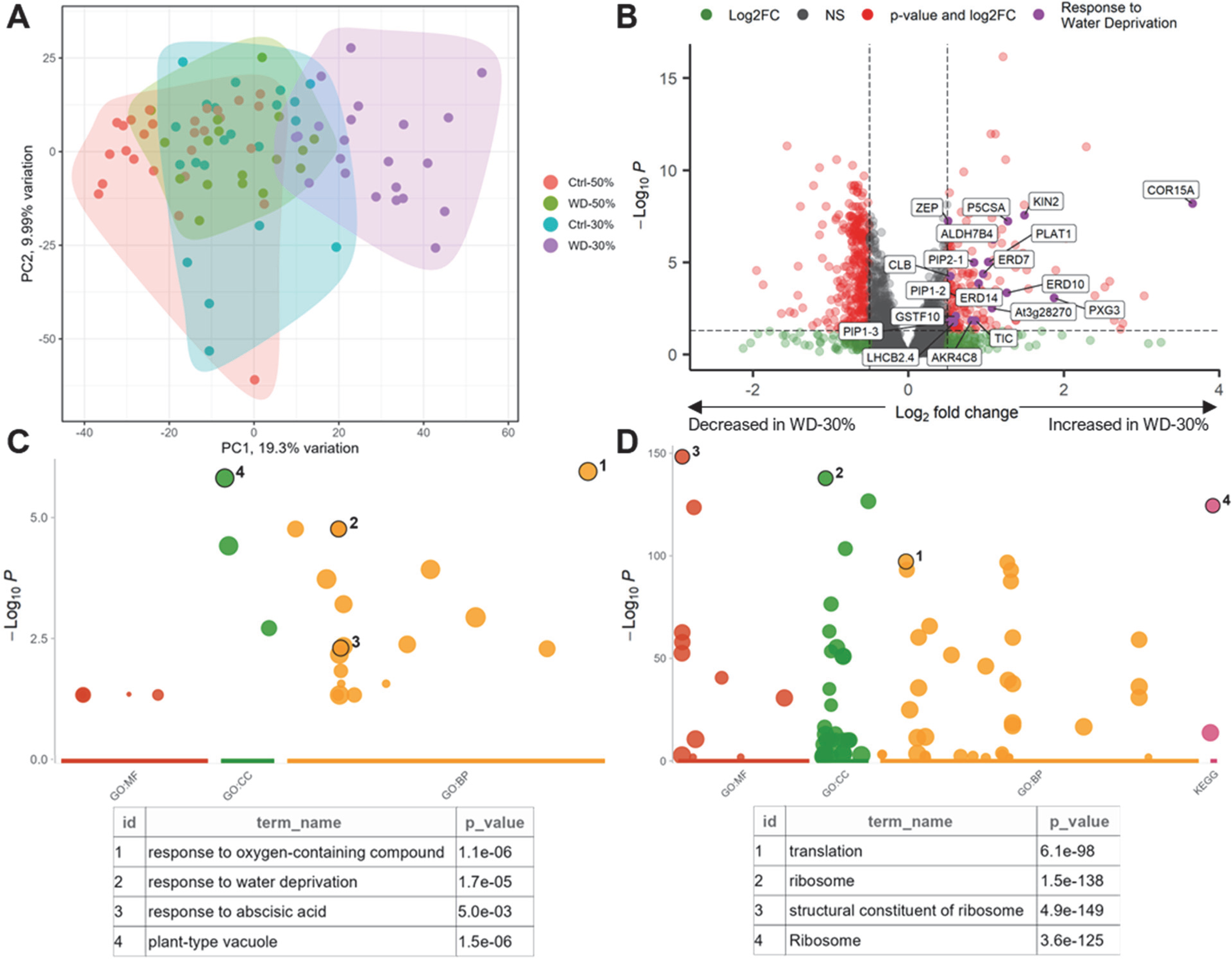
Changes in protein abundance under severe water deficit stress revealed by nanoPOTS scProteomic analysis. **(A)** Principal component analysis of control and water deficit protoplasts. (**B**) Differential abundance analysis of Ctrl-30% and WD-30% protoplasts, highlighting statistically significant proteins. Labeled proteins were annotated by the GO term “response to water deprivation”. Dashed lines indicate log_2_ fold-change (≥ 0.5 and ≤ -0.5) and BH adjusted p-value thresholds ( ≤ 0.05) for considering a change in abundance as significant. (**C**) Manhattan plot of statistically significant GO terms (BH adjusted p-value ≤ 0.05) associated with proteins increasing in abundance. Color indicates GO domain. Selected terms are circled and presented in tables below. (**D**) Same as (**C**) but with statistically significant proteins decreasing in abundance.

Although limited separation was achieved with WD-50% compared to control, WD-30% provided more apparent separation from Ctrl-30%. Comparison of protein abundances revealed few statistically significant differences in the context of WD-50%, with only 28 proteins passing the log2FC cutoff of 0.5/-0.5 and FDR of < 0.05 (supplemental Fig. S10 and supplemental table S2). On the other hand, WD-30% showed many more statistically significant differences in protein abundance with 169 upregulated proteins and 294 downregulated proteins (Fig. 3B and supplemental table S2), which is also larger than previously reported by bulk shotgun proteomics ^21^. This can likely be attributed a few factors, specifically: (1) The greater homogeneity of our sample, which is enriched for a single-cell type and (2) the higher sensitivity methodology adopted by our scProteomics workflow. Of the upregulated proteins, 19 are known to be involved in response to water deprivation (Fig. 3B).

Gene ontological (GO) and KEGG enrichment of proteins significantly (BH adjusted p-value < 0.05) increased (log2FC ≥ 0.5) or decreased (log2FC ≤ -0.5) in abundance in the WD-30% group revealed a number of term enrichments. Notably, among proteins increasing in abundance under water-deficit conditions were terms related to drought stress pathways, including “response to water deprivation”, “response to abscisic acid”, and “water channel activity” (Fig. 3C and supplemental table S3). For proteins decreasing in abundance, enriched terms including “ribosome” and “translation” were observed with p-values exceeding 1 x 10^-90^ (Fig. 3D and supplemental table S4). This level of significance can be attributed to the observation that over 50% of proteins decreasing in abundance (160 proteins) were considered “structural constituents of the ribosome”, indicating a significant loss of translational capacity in WD-30% plant leaf mesophylls. It is worth noting our results align well with Heinemann et al. who similarly found strong abundance decreases among ribosomal proteins^21^.

### Covariation analysis reveals protein-protein dynamics under water deficit stress

We next looked beyond differential abundance testing and leverage our single-cell data to investigate protein covariation in the context of water deficit stress.

Considering protein covariation infers functional or physical protein interactions, we wondered if our data might provide insights into protein-protein interactions or changes in protein covariation occurring independently of changes in differential abundance in response to drought stress. Towards this end, we subset proteins by applying a conservative cut-off of >90% observations for each protein across all 87 protoplasts, imputed with KNN imputation for the remaining missing observations, and combined GO enrichment analysis with GMM clustering (k = 19) to identify covariant protein clusters.

After GMM clustering 2,323 proteins were assembled into 19 clusters, 18 of which had statistically significant GO term enrichments (see supplemental table S5 for all enriched terms). In cases where multiple GO terms were assigned to the same cluster, the term with the most significant p-value was chosen as the representative term. To visualize this consensus protein network, we calculated all pairwise Pearson correlations and mapped these correlations to the 18 clusters with statistically significant GO terms (Fig. 4A and supplemental table S6).

**Fig. 4.**
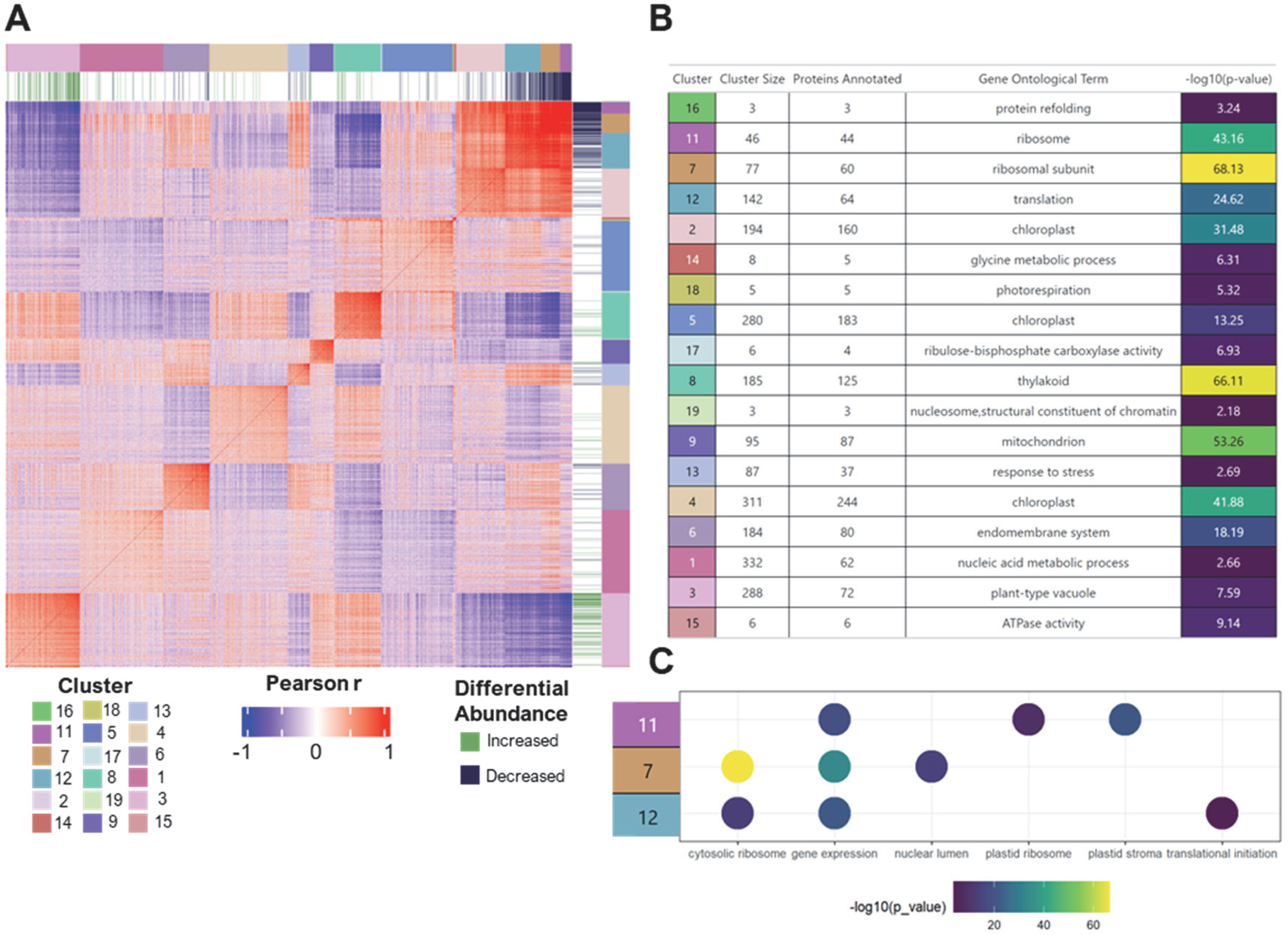
GMM clustering guided by GO enrichment accurately identifies functional protein networks. **(A)** Symmetrical matrix of pairwise Pearson correlations for 2,323 proteins organized into 18 clusters represented with at least one statistically significant GO term. Annotations alongside the matrix indicate if a protein was found to be differentially abundant under severe water deficit stress (WD-30%), as well as the cluster each protein belongs to. (**B**) Cluster descriptive statistics and representative GO terms for each cluster. -log_10_ BH adjusted p values are indicated by color as well. (**C**) Selected GO terms and their statistical significance for clusters 11, 7, and 12.

Within the 18 clusters, we observed enrichment for many cellular components across diverse molecular functions (Fig. 4B). Clusters ranged in membership from over 300 proteins (cluster 4) down to just 3 proteins (cluster 19), highlighting the strength of our approach in clustering across a large dynamic range. Chloroplast, the most abundant organelle in mesophyll cells, was represented in three clusters (clusters 2, 5, and 4) as demonstrated with the representative GO term “chloroplasts”. A separate cluster was also found to have distinct enrichment for the thylakoid compartment of chloroplasts (cluster 8), with 125 out of 185 proteins annotated as such within the cluster (Fig. 4B). Although these clusters are part of the same organelle, they display very different correlation patterns across the network, which demonstrates these clusters are not just a consequence of overfitting or cluster splitting with the GMM clustering approach. Indeed, it can be observed that the thylakoid cluster (cluster 9) shows a trend toward negative correlations with the chloroplast-related cluster 2 (Fig. 4A). Clusters 11, 7 and 12 demonstrated the highest within-cluster correlations and were all enriched in ribosomal proteins, with many enriched terms reflecting involvement in translation and gene expression (Fig. 4C). These clusters also showed notable differences with cluster 11 being more closely related to chloroplast ribosomal complexes, cluster 7 showing larger enrichment for cytosolic ribosomal proteins, and cluster 12 being more represented in translation initiation (Fig. 4C). Cluster 11 is also notable in that it contained a single protein within it that was not associated with the term “ribosome” in the GO database (AT5G14910 or CRASS, supplemental Fig. S11). This single protein, despite being annotated as a heavy metal transport protein, was only recently found to be potentially associated to the ribosome based on co-immunoprecipitation experiments ^43^. This study showed CRASS interact with plastid ribosomal proteins PRPS1 (AT3G13120) and PRPS5 (AT2G33800), which were both found within cluster 11 and strongly correlated with CRASS (supplemental Fig. S11).

Collectively, these data demonstrate scProteomics-enabled covariation analysis holds great potential to reveal previously unrecognized protein functions and protein-protein interactions by clustering them together within functional modules. This capability is particularly useful for plant systems whose gene annotation information is lacking for many proteins or inferred mainly from orthologues.

To facilitate a better understanding of the changes in protein regulatory dynamics in the context of severe water deficit stress, we separately calculated the mean Pearson correlations for Ctrl-30% (n = 21) and WD-30% (n= 22) for each cluster. Correlations from Ctrl-30% were then subtracted from WD-30% to provide the matrix of correlation differences. Based on changes in average correlation across groups, we could observe several clusters with proteins that had striking shifts in their overall correlation distributions (Fig. 5A; full, unaveraged heatmaps are shown in supplemental Fig. S12).

**Fig. 5.**
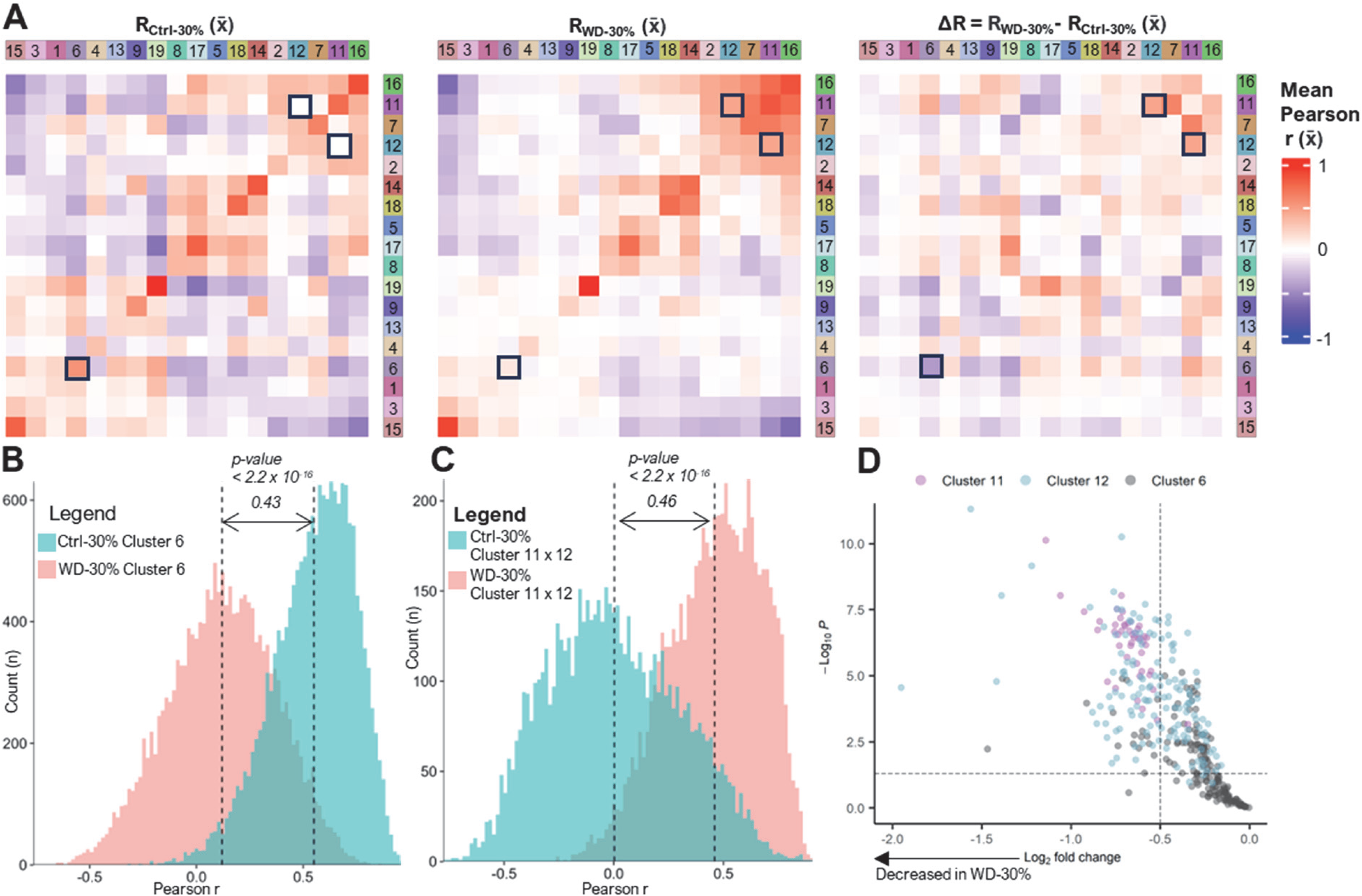
Dynamic changes in protein covariation due to water deficit stress captured by nanoPOTS scProteomics. **(A)** Mean (x̄) Pearson correlation matrices for Ctrl-30%, WD-30%, and the matrix of correlation differences (ΔR) organized by cluster. Black boxes indicate cluster 6 and the intersection between clusters 11 and 12. Note, for visualization purposes, clusters are not scaled by the number of proteins within each cluster and instead displayed equally in size. (**B**) Distributions of cluster 6 correlations for control (Ctrl-30%, blue) and severe water deficit (WD-30%, magenta) protoplasts. The means of each distribution are represented with dashed lines, with the difference shown between them. Significance was determined using the Kolmogorov-Smirnov test. (**C**) Same as (**B**) but with the intersection between clusters 11 and 12. (**D**) Differential abundances of proteins found in clusters 11, 12, and 6. Dashed lines indicate log_2_ fold-change (≤ -0.5) and BH adjusted p-value thresholds ( ≤ 0.05) for considering a change in abundance as significant.

Based on changes in average correlation across groups, we could observe several clusters with proteins that had striking shifts in their overall correlation distributions. For example, over one third of cluster 6’s (endomembrane system) pairwise correlations (6,282) were considered statistically significant (FDR < 0.05) under no water stress.

However, introduction of severe water stress led to a near complete loss of this covariation pattern, with only 178 pairwise correlations being significant and the entire distribution shifting to a mean correlation near 0 (Fig. 5B). A similar impact was seen between clusters 11 (ribosome) and 12 (translation), however in this case severe water stress induced stronger correlations between these two clusters (Fig. 5C). These two examples also uniquely highlight how protein covariation can be both related and unrelated to changes in mean protein abundance. Ribosomal proteins are strongly decreased in abundance, including those in clusters 11 and 12 (Fig. 5D). On the other hand, the endomembrane system (cluster 6) suggests differential abundance may not fully capture underlying changes in protein-protein relationships as few proteins (16 out of 184) showed statistically significant change in abundance despite strong changes in the underlying protein covariation (Fig. 5D).

## Discussion

Our work represents the first application of label free single-cell proteomics to plant systems, established with the model organism *Arabidopsis thaliana*. Using the tape-sandwich protoplasting approach, piezoelectric cellenONE sorting, and nanoPOTS techniques, we demonstrated high-quality mesophyll protoplast isolation with downstream single-cell proteomics analyses. As mesophyll cells do not appear to have unique cell type markers ^44^, we relied on the observation that they instead are enriched for proteins involved in photosynthesis (LHCs) and carbon fixation (RuBisCo). Indeed, it has been demonstrated that the fractional volume occupancy of chloroplasts in mesophylls is much higher (9.3% for mesophyll palisade and 15.5% for spongy mesophyll) than epidermal cells (0.45% for abaxial and 0.22% for adaxial)^45^. In agreement with the high-volume chloroplast occupancy in the mesophyll cell type, we observe LHC and RuBisCo proteins represent up to 46% of protein copies on average in well-watered protoplasts in this study. The high abundance of RuBisCo and LHC proteins, as well as strong overlap in protein abundances with prior proteomic analysis on bulk Arabidopsis leaf tissue ^21^, clearly demonstrate enrichment of the mesophyll cell type in our study.

Precision, accuracy, and quality of single-protoplasts were assessed with standardized metrics and agreed with prior label free single-cell studies in cultured mammalian cells^46–48^. We found protoplast fragmentation could have strong negative effects on scProteomic data quality but could be mitigated by the cellenONE system with low-pressure cell sorting and imaging capabilities. Presumably, protoplast fragmentation is caused by loss of the cell wall, as removal of the cell wall is known to increase the fragility and reduced integrity of these cells^25^. Therefore, we suggest caution to future experimentalists performing single-protoplast proteomics but also believe our approach can serve as a useful guide to isolate high-quality protoplasts for downstream analysis.

One of the main advantages of single-cell proteomics is that it can provide context-dependent insights into protein-protein relationships on a global scale without the need for intermediate affinity reagents or for exogenous expression of tagged baits ^49–52^. Protein covariation across large-scale proteomic studies are implicit indicators of protein-protein relationships^53–55^. However, the averaging of protein measurements in bulk proteomic analyses weakens these relationships. Hence, more direct and contextual protein-protein association information can only be obtained when protein covariation measurements are acquired at the single-cell level. Protein pairs that covary may reflect direct interactions (such as subunits within a complex) or functionally related proteins (such as enzymes that process substrates in a metabolic pathway). To this end, we have described a generalizable approach for constructing functional protein networks at the single-cell level using GMM clustering guided by known gene ontologies. Application of our pipeline to 87 Arabidopsis protoplasts in the context of water deficit stress provided 18 protein clusters, several of which partially or completely comprised proteins from known complexes such as the glycine cleavage system (cluster 14), vacuolar ATP synthase (cluster 15), chaperonin 60 (cluster 16), RuBisCo (cluster 17), and several ribosomal assemblies. Importantly, several clusters showed changes in protein covariation without significant changes in abundance, highlighting how scProteomics can reveal dynamics that may be missed by shotgun proteomic approaches which often only characterize differential abundance. Furthermore, the assembly of covarying proteins into distinct clusters allowed us to identify novel protein functions using a “guilt by association” approach^56^. We observed a protein (CRASS) annotated as a heavy metal transport protein tightly correlated with a cluster of chloroplast ribosome proteins. Interestingly CRASS was only recently found to be potentially associated with the ribosome^43^. Our data validates the identification of CRASS as a chloroplast ribosomal protein and highlights how clustering can be used to identify new protein relationships.

Few statistically significant differences were found in plants exposed to a moderate water deficit (WD-50%) compared to those that were well watered.

Retrospectively, we believe this may be due to these plants being younger and perhaps less responsive to the relatively moderate water deprivation. Indeed, plant age and abiotic stress response are clearly linked, however this is still an area of active research^57^. In the context of severe water deficit stress (WD-30%), we observed a striking reduction in abundance of 160 ribosomal proteins, substantiating prior work by Heinemann et al. who identified 125 ribosomal proteins with losses in abundance in response to increasing water deficit stress^21^. Considering ribosomes account for a large proportion of a cell’s RNA and protein content we believe our results support a process where increased turnover of ribosomes under stress conditions are a mechanism for allocating amino acid and nucleic acid building blocks to different plant tissues, cell types, and/or pathways^21,58^. Surprisingly, the protein covariation data also demonstrated that decreases in ribosomal protein abundance were inversely related to increases in protein-protein correlations for some ribosome and translation related proteins.

One interpretation of this result is that the mechanisms that are driving ribosome degradation are selective for subunits that exceed the necessary stoichiometry to form active complexes. In other words, the overproduction of a given protein may be tolerated under homeostatic conditions but not when under water-deficit stress.

Interestingly, turnover of entire ribosome assemblies through ribophagy is a known abiotic stress response mechanism in plants ^21,59,60^. While ribophagy is a demonstrated response to stress in plants, our covariation data also suggests alternative pathways with greater selectivity such as the Excess RIboSomal protein Quality control (ERISQ) pathway must also play a role in plant stress response^61,62^. Ribophagy as a mechanism for amino acid allocation is also linked to processes involved in proline synthesis as plants utilize proline as an osmolyte protectant when exposed to abiotic stresses ^63,64^.

Indeed, we found the rate-limiting enzyme in the proline synthesis pathway (delta-1-pyrroline-5-carboxylate synthase A, PGCSA or PGCS1) to be increased in abundance by 2.4-fold under severe water-deficit stress while the non-stress responsive P5CS2 ^65^ was unchanged. Plants within stressed environments can also utilize amino acids as an alternative for mitochondrial respiration substrates during inadequate carbohydrate supply due to a decrease in photosynthesis rates ^66^. The degradation pathways for Lys and branched-chain amino acids such as Val, Leu, and Ile have previously been identified as crucial factors for *Arabidopsis* dehydration tolerance ^67^. Furthermore, aromatic amino acids produce a wide range of secondary metabolites, and we observed proteins involved in secondary metabolism pathways (such as GA2OX6, ABA1, CAT2/3, AMY3, BAM2/6, and ACO3) to have an increase in abundance in response to severe water deficit stress.

## Conclusions

Abiotic stresses such as drought are increasingly impacting crops worldwide and new approaches for understanding stress-response at the single-cell level are needed. The label-free scProteomic approach presented here represents a significant advance through the demonstration of a facile protoplast isolation method combined with deep and precise proteomic coverage of Arabidopsis leaf mesophyll cell types. We have also described a computational approach for analyzing protein covariation patterns, one of the main but underutilized advantages of performing scProteomics. Applying the full methodology to water-stressed plants provided data that not only aligned with prior studies but has orthogonally validated only recently discovered results and generated new hypothesis for future studies of plant responses to abiotic stresses. In summary, we believe the methodology and results presented here will serve as an informative reference to future scProteomic studies as well as investigations into abiotic stresses.

## Data Availability

All scripts, functions, and source data are available at the Github repository https://github.com/Cajun-data/nanoPOTS_Arabidopsis. The mass spectrometry raw data have been deposited to the ProteomeXchange Consortium via the MassIVE partner repository with dataset identifier MSV000095072.

## Acknowledgements

We thank Dr. Matthew Monroe for his assistance in depositing the raw proteomic data onto MassIVE. This work was supported by a Laboratory Directed Research and Development award (I3T) from Pacific Northwest National Laboratory (Y.Z.) and project award (10.46936/intm.proj.2020.51688/60000255, Y.Z.) from the Environmental Molecular Sciences Laboratory, a DOE Office of Science User Facility sponsored by the Biological and Environmental Research program under Contract No. DE-AC05-76RL01830.

## CRediT authorship contribution statement

James M. Fulcher: Conceptualization; Investigation; Validation; Methodology; Formal Analysis; Writing – Original Draft, Reviewing, and Editing; Visualization. Pranav Dawar: Formal Analysis; Visualization; Writing –Reviewing and Editing. Vimal Kumar Balasubramanian: Conceptualization; Methodology; Formal Analysis; Investigation; Writing – Reviewing and Editing. Sarah M. Williams: Investigation; Writing – Reviewing and Editing. Kyle D. Nadeau: Software; Writing – Reviewing and Editing Tanya E. Winkler: Investigation; Writing – Reviewing and Editing. Lye Meng Markillie: Methodology; Investigation; Writing – Reviewing and Editing. Hugh D. Mitchell: Data Curation; Writing – Reviewing and Editing. Amir H. Ahkami: Conceptualization; Writing – Reviewing and Editing; Funding acquisition. Ljiljana Paša-Tolić: Supervision; Project administration; Writing – Reviewing and Editing. Ying Zhu: Supervision; Conceptualization; Methodology; Investigation; Project administration; Funding acquisition; Writing – Reviewing and Editing.

## Abbreviations

FAIMS: high Field Asymmetric waveform Ion Mobility Spectrometry
scProteomics: single-cell Proteomics
TIFF: Transferring Identifications based on FAIMS Filtering
LFQ: label free quantification
nanoPOTS: nanodroplet Processing in One Pot for Trace Samples
SWC: soil water content
WD: water deficit
AGC: Automatic Gain Control
GO: Gene Ontology
CV: Coefficient of Variation
DDM: n-Dodecyl-β-D-Maltoside
IAA: Iodoacetamide
ABC: Ammonium Bicarbonate
FA: Formic Acid
PEEK: Polyetheretherketone
DTT: Dithiothreitol
PFDS: heptadecafluoro-1,1,2,2-tetrahydrodecyl-dimethylchlorosilane
FDR: false discovery rate
HCD: higher-energy collisional dissociation)

