## Supplementary Information for "Single-cell proteomics of Arabidopsis leaf mesophyll identifies drought stress-related proteins"

**Table of Contents:**


| Title | Page |
|---|---|
| Figure S1– Arabidopsis growth conditions and overview | 3 |
| Figure S2– Assessment of data completeness and sensitivity | 4 |
| Figure S3– Images of tape-sandwich approach | 5 |
| Figure S4– Pearson correlations for scProteomics of 10, 3, and 1 protoplast | 6 |
| Figure S5– cellenONE images of protoplasts during sorting | 7 |
| Figure S6– PCA analysis of control and water deficit protoplasts | 8 |
| Figure S7– Clustering, comparison, and images of fragmented protoplasts | 9 |
| Figure S8– Abundance comparison of LHC proteins | 10 |
| Figure S9– Comparisons with Heinemann et al 2012 | 11 |
| Figure S10– Volcano plot of moderate water deficit protoplasts | 12 |
| Figure S11– Heatmap of ribosomal proteins from cluster 11 | 13 |
| Figure S12– Full Pearson correlation matrices | 14 |

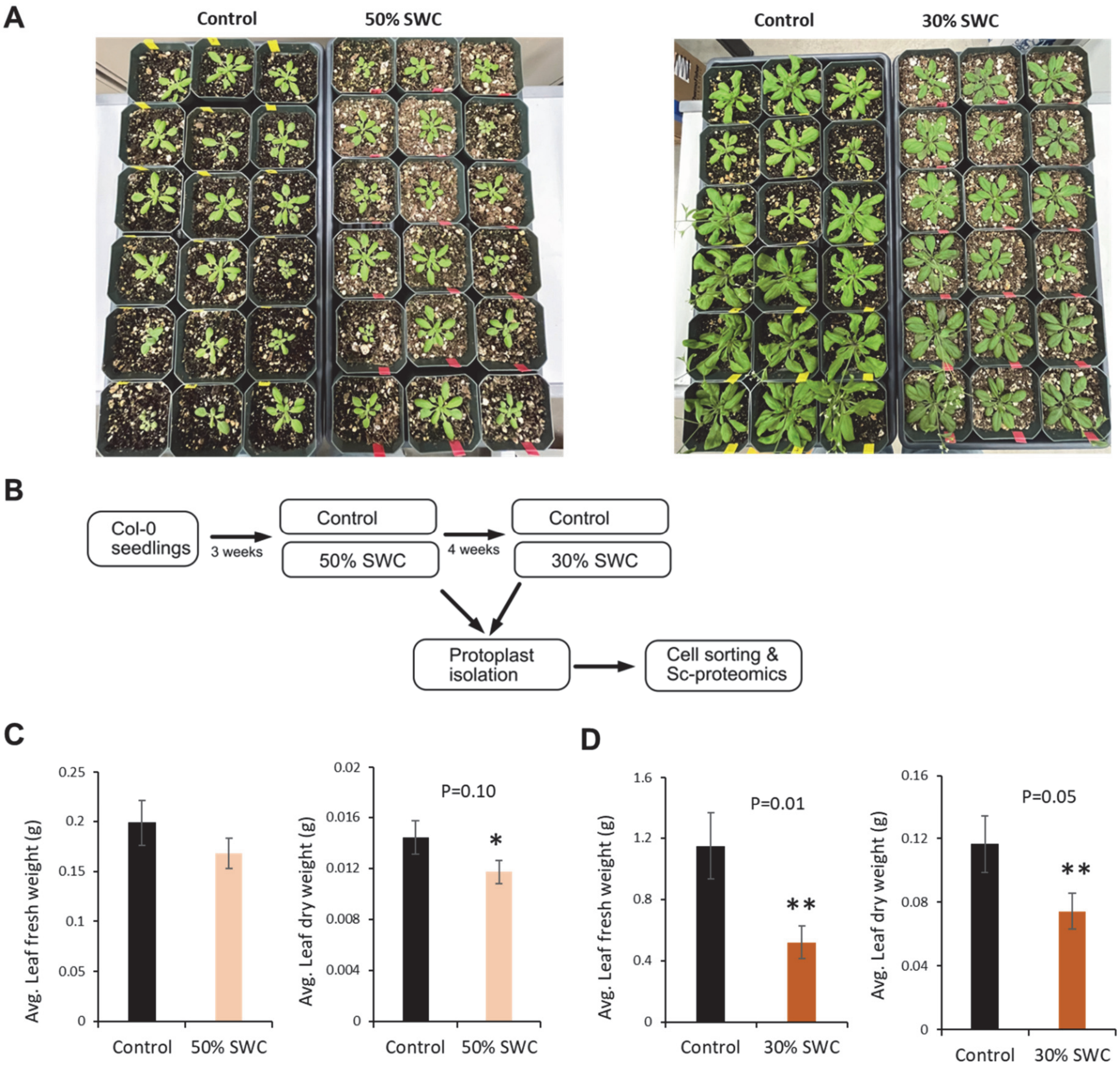


**Figure S1** (**A)** Trays of Arabidopsis Col-0 ecotype plants that include well-watered (control) plants and water deficit (WD) stress treated plants by adjusting soil water content (SWC) to 50% and 30% of corresponding control pot weights. (**B**) Workflow describing the timeline involved in creating 50% and 30% SWC stress treatments and samples collected for protoplasting and scProteomic analysis. (**C**) Average fresh and dry weights of leaf tissue obtained from control and WD (50% SWC and 30% SWC) stress exposed Col-0 plants.

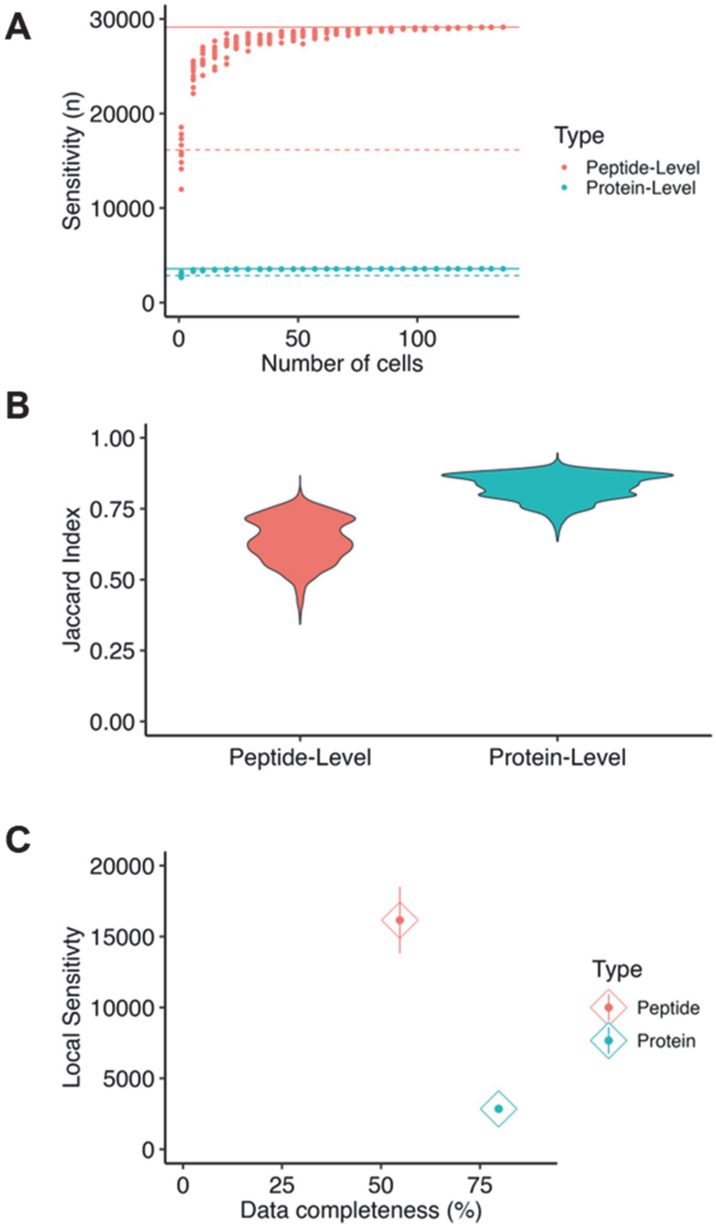


**Figure S2** (**A**) Cumulative sensitivity curve showing the total number of distinct peptides and proteins (as indicated by color) as an increasing number of single protoplasts are sampled (n = 136 total). The dashed lines represent local sensitivity while the solid lines represent total sensitivity. (**B**) Distributions of the pairwise Jaccard indices at the peptide and protein level. (**C**) Average local sensitivity with respect to data completeness at the peptide and protein level.

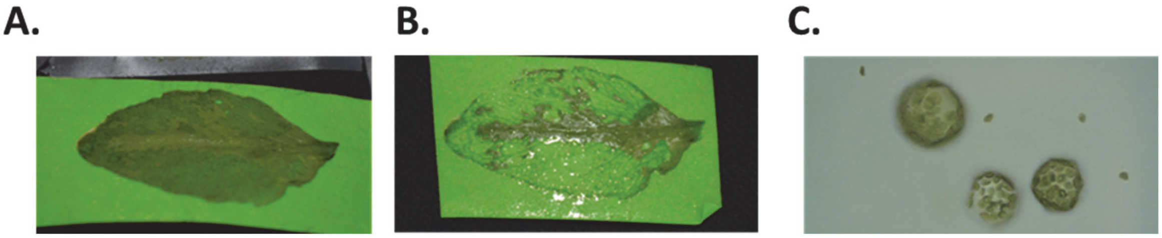

**Figure S3** (**A**). Fully grown rosette leaf used for tape sandwich method where the abaxial side was peeled to expose the inner adaxial side with enriched palisade mesophyll cells. (**B**). Cell wall digestion was performed using protoplast isolation buffer leaving the remains of vascular cells enriched midveins onto the tape (**C**). Isolated protoplasts (size ~30-50$\mu$m) count was around one million protoplasts/ml of buffer.

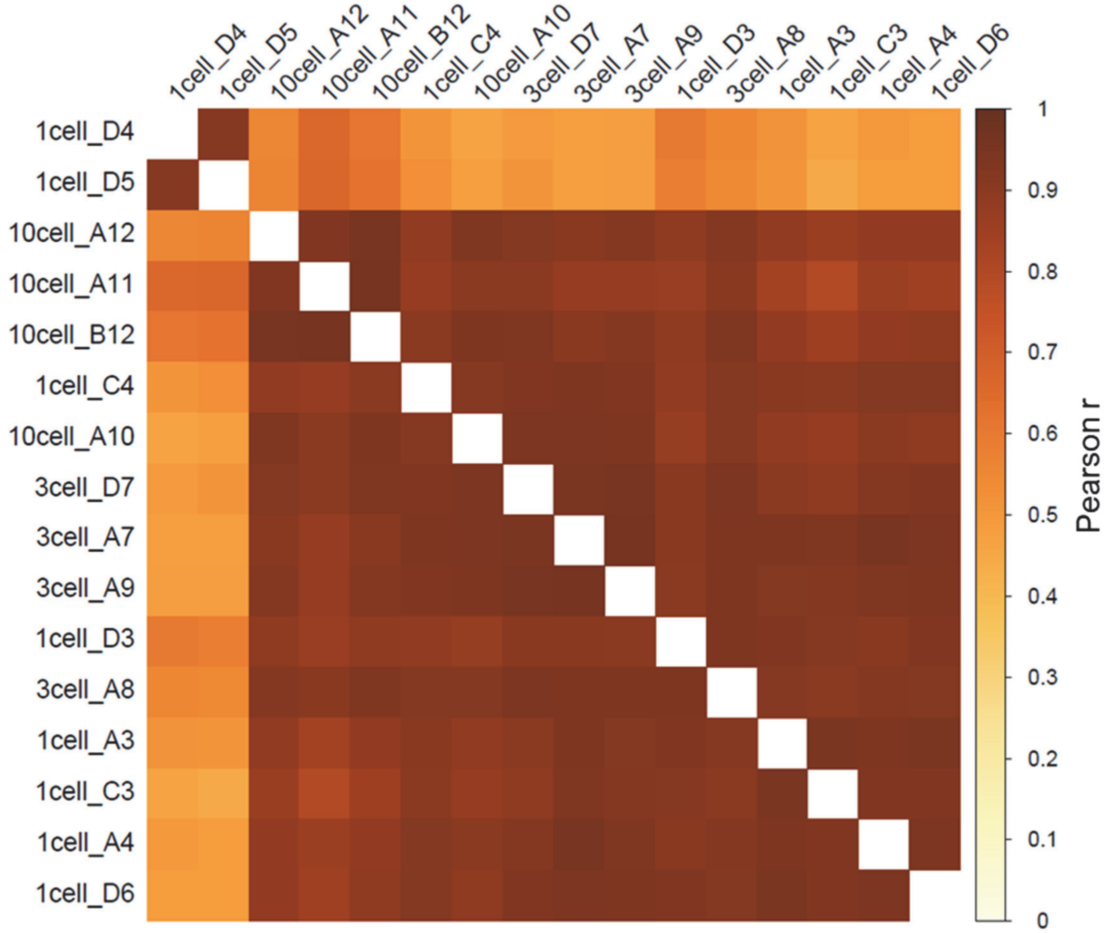


**Figure S4** Heatmap of pairwise Pearson correlations based on $log_2$ protein abundances.

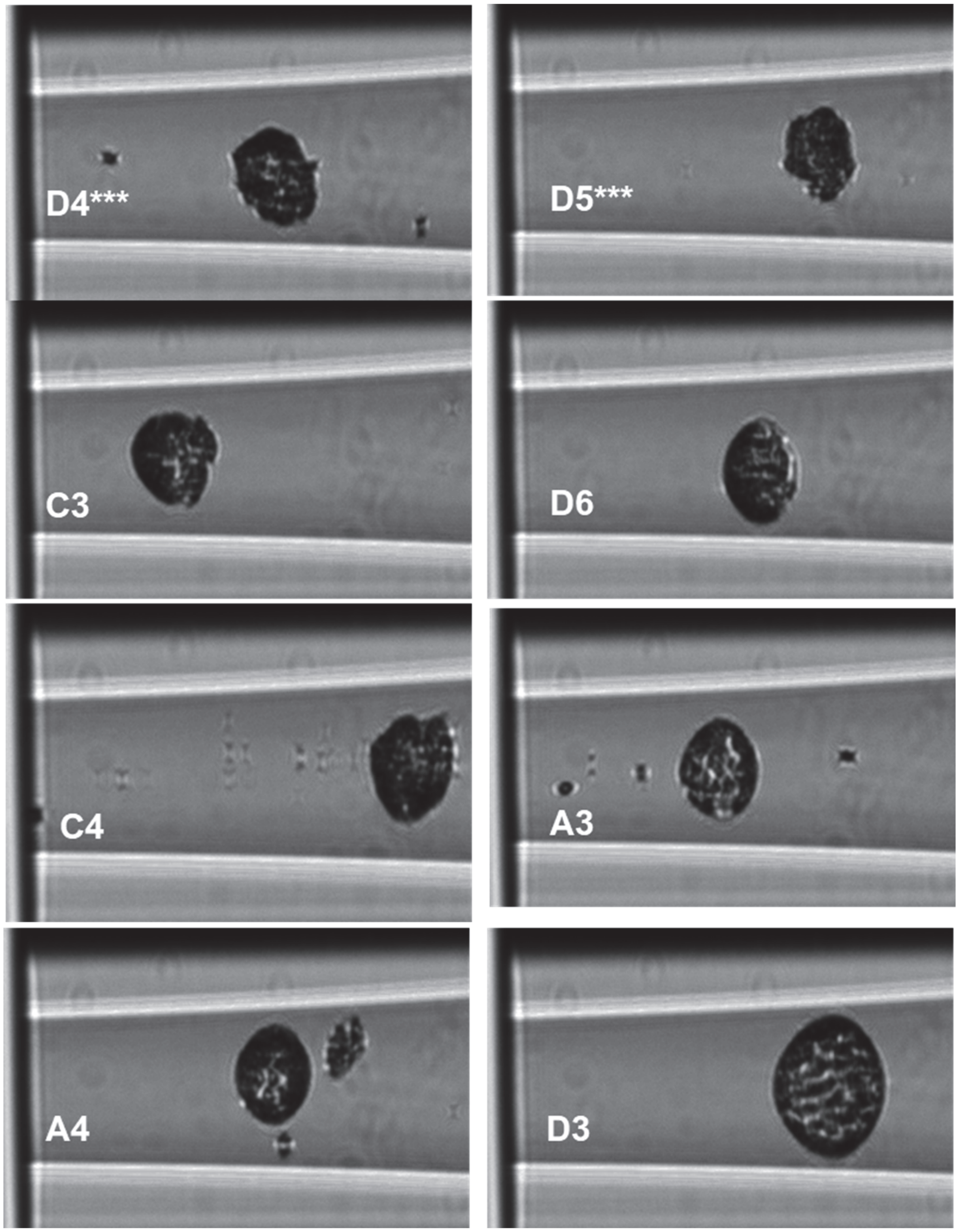


**Figure S5** Images taken of sorted single protoplasts during cell sorting on the cellenONE. Sample names (indicating well position on the nanoPOTS chip) are shown in the bottom left corner. *** represents outlier protoplasts identified in **Figure S4**.

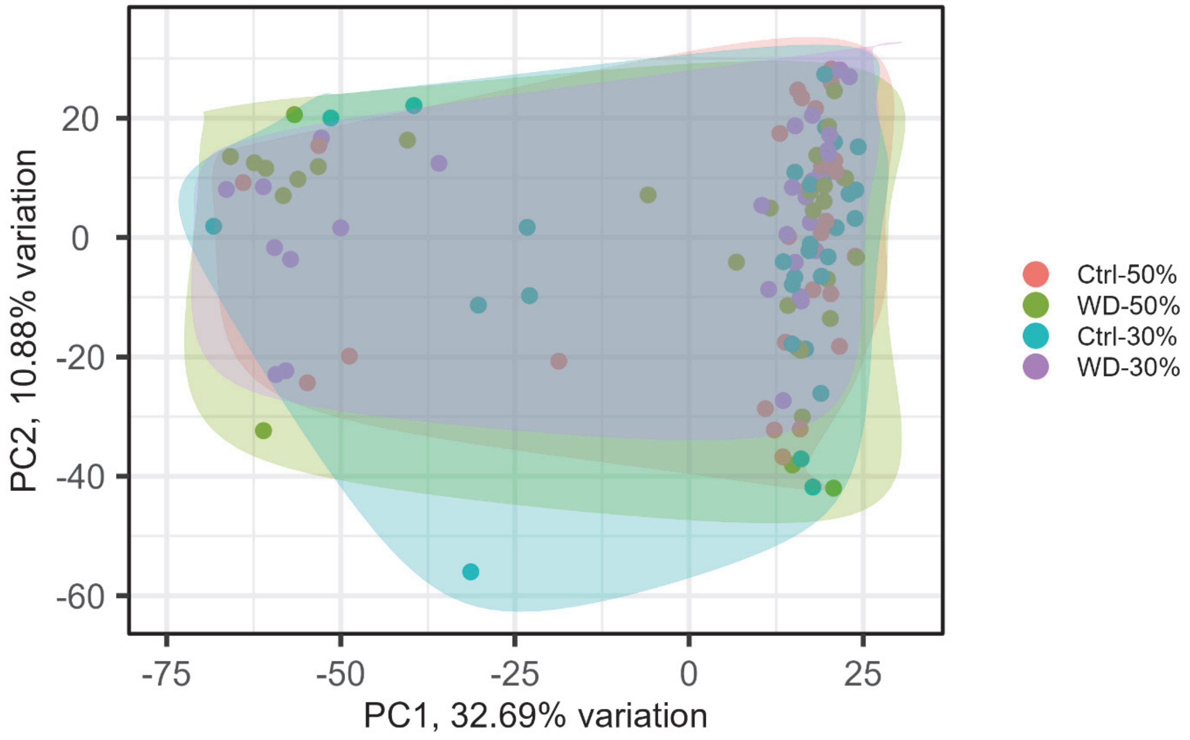


**Figure S6** PCA of all 117 protoplasts across two control groups (Ctrl-50% and Ctrl-30%) and two water deficit groups (WD-50% and WD-30%).

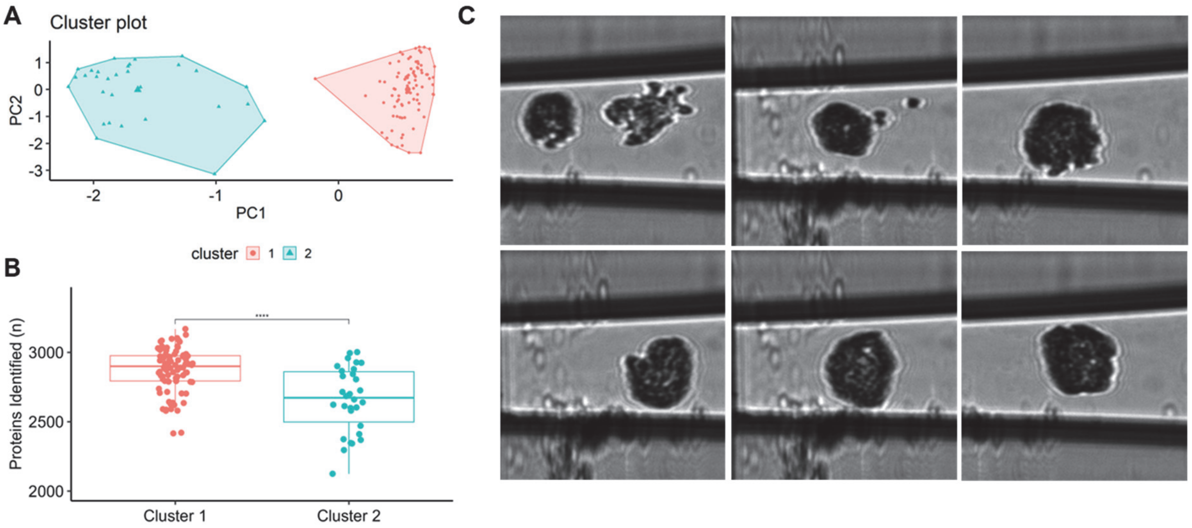


**Figure S7** (**A**) PCA of all 117 protoplasts with kmeans clustering (k = 2). (**B**) Box plots showing the number of protein identifications (y-axis) for protoplasts falling cluster 1 or cluster 2. Statistical significance was determined using t-test (**** indicates p value ≤ 0.0001). (**C)** Representative cellenONE images of fragmented protoplasts belonging to cluster 2.

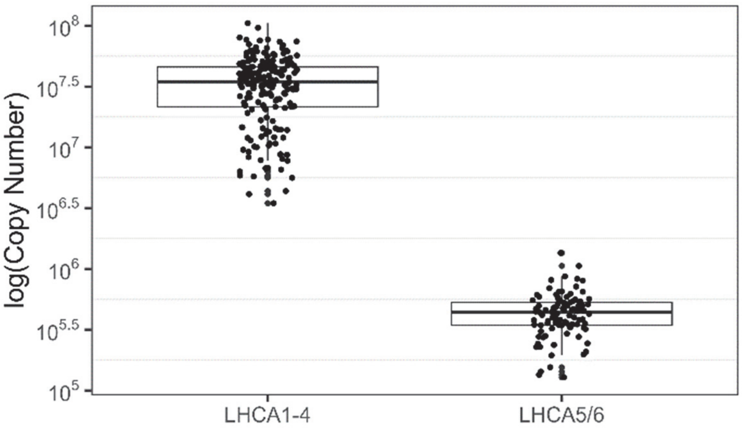


**Figure S8** Box plot comparing the $log_{10}$(Copy Number) of LHCA(1-4) and LHCA(5-6) across protoplasts

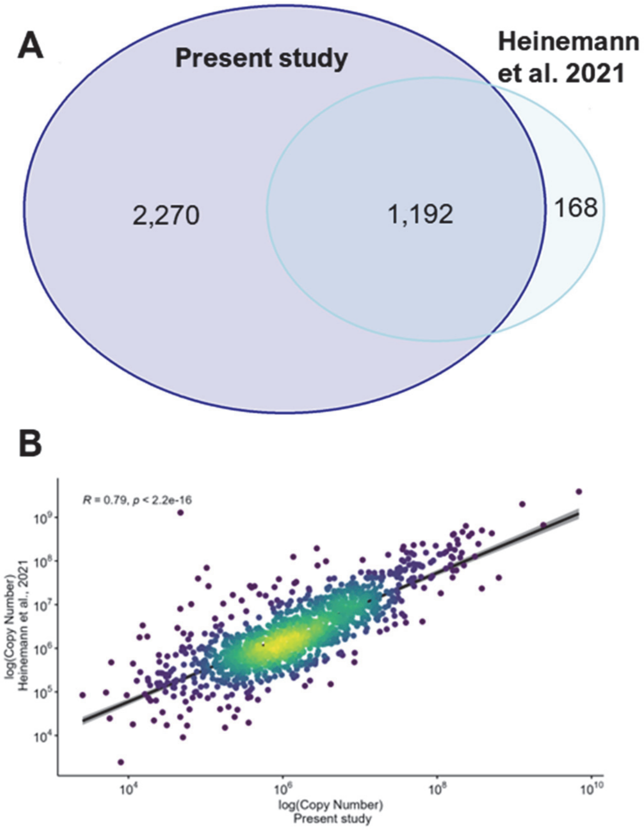


**Figure S9** (**A**) Venn diagram of proteins identified in this study and Heinemann et al. 2021. (**B**) Scatterplot of log-transformed protein copy numbers from Heinemann et al. 2021 and this study. Linear regression is shown with confidence intervals. Pearson correlation is shown in upper left corner.

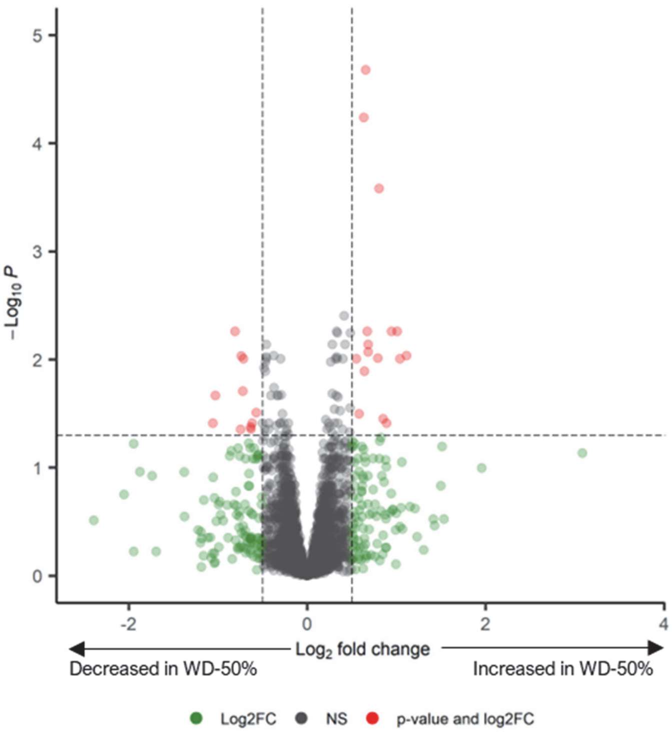


**Figure S10** Volcano plot comparing WD-50% and Ctrl-50% protein abundances. Dashed lines indicate $\log_2$ fold-change (≥ 0.5 and ≤ -0.5) and adjusted p-value cutoffs ( ≤  0.05)

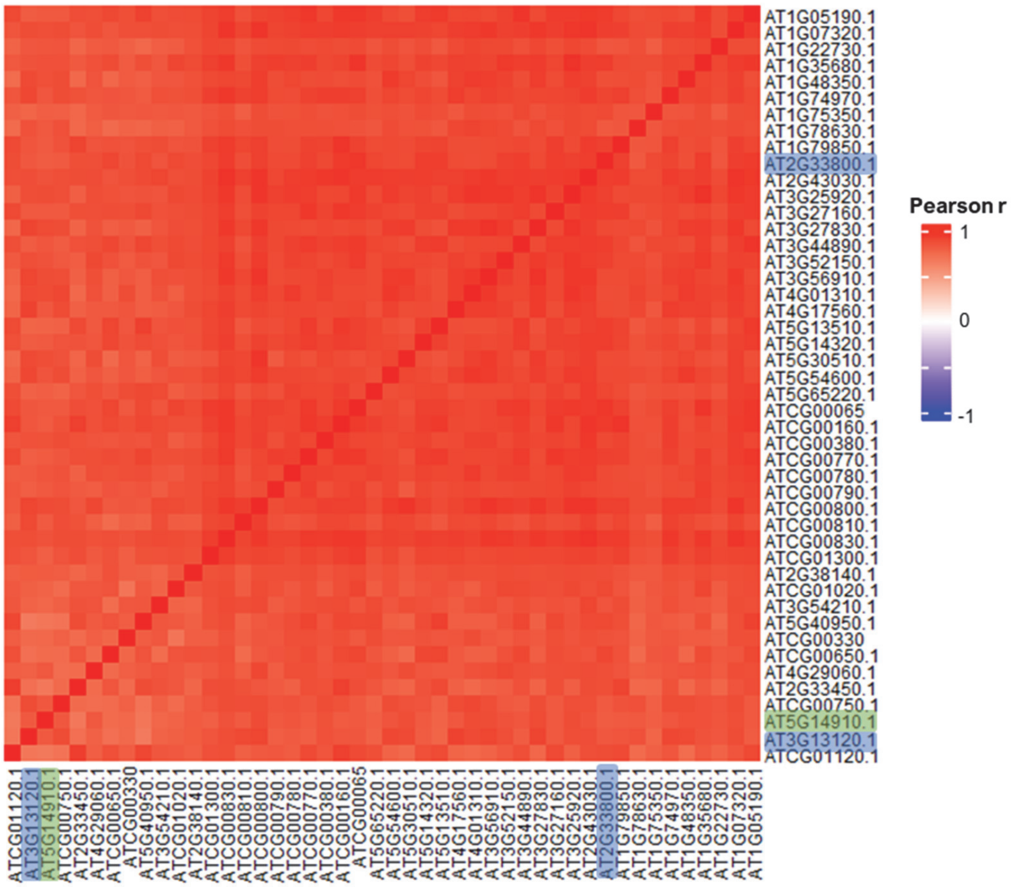


**Figure S11** Heatmap of Pearson correlations for ribosomal proteins within cluster 11 (46 proteins total). Green shading of indicates AT5G14910 which lacks GO annotation for "ribosome" term. Blue shading indicates known interacting partners AT5G14910

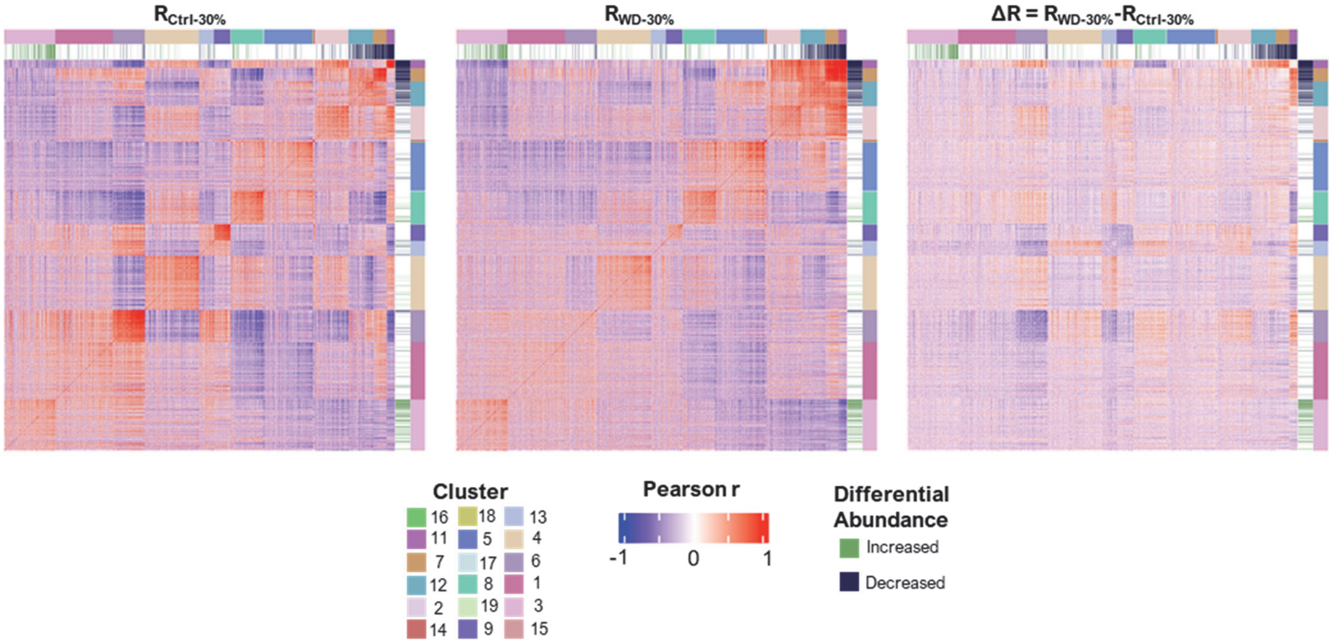


**Figure S12** Full correlation matrices (left $R_{Ctrl-30\%}$, middle $R_{WD-30\%}$, and right ΔR) consisting of 2,252 proteins arranged by Gaussian Mixture Modeling clustering (k = 19). Annotations directly next to matrices indicate proteins found to be differentially abundant ($log_2FC \geq 0.5$ or $\leq -0.5$ with adjusted p value $< 0.05$) while the second row of annotations indicates the cluster each protein belongs to. Matrix ΔR (right) represents the difference in correlation between the two groups.
